# Design of Specific Primer Sets for the Detection of SARS-CoV-2 Variants of Concern B.1.1.7, B.1.351, P.1, B.1.617.2 using Artificial Intelligence

**DOI:** 10.1101/2021.01.20.427043

**Authors:** Carmina A. Perez-Romero, Alberto Tonda, Lucero Mendoza-Maldonado, Etienne Coz, Patrick Tabeling, Jessica Vanhomwegen, Eric Claassen, Johan Garssen, Aletta D. Kraneveld, Alejandro Lopez-Rincon

**Affiliations:** Division of Pharmacology, Utrecht Institute for Pharmaceutical Sciences, Faculty of Science, Utrecht University, Universiteitsweg 99, 3584 CG Utrecht, the Netherlands; Departamento de Investigación, Universidad Central de Queretaro (UNICEQ), Av. 5 de Febrero 1602, San Pablo, 76130 Santiago de Querétaro, Qro., Mexico; UMR 518 MIA-Paris, INRAE, c/o 113 rue Nationale, 75103, Paris, France; Hospital Civil de Guadalajara “Dr. Juan I. Menchaca”. Salvador Quevedo y Zubieta 750, Independencia Oriente, C.P. 44340 Guadalajara, Jalisco, México; Athena Institute, Vrije Universiteit, De Boelelaan 1085, 1081 HV Amsterdam, the Netherlands; Department Immunology, Danone Nutricia research, Uppsalalaan 12, 3584 CT Utrecht, the Netherlands; Julius Center for Health Sciences and Primary Care, University Medical Center Utrecht, Universiteitsweg 100, 3584 CG Utrecht; CBI, ESPCI Paris, Université PSL, CNRS, 75005 Paris, France; CIBU, Institut Pasteur, 25-28 rue du Dr Roux, Paris 15

## Abstract

As the COVID-19 pandemic continues, new SARS-CoV-2 variants with potentially dangerous features have been identified by the scientific community. Variant B.1.1.7 lineage clade GR from Global Initiative on Sharing All Influenza Data (GISAID) was first detected in the UK, and it appears to possess an increased transmissibility. At the same time, South African authorities reported variant B.1.351, that shares several mutations with B.1.1.7, and might also present high transmissibility. Earlier this year, a variant labelled P.1 with 17 non-synonymous mutations was detected in Brazil. Recently the World Health Organization has raised concern for the variants B.1.617.2 mainly detected in India but now exported worldwide. It is paramount to rapidly develop specific molecular tests to uniquely identify new variants. Using a completely automated pipeline built around deep learning and evolutionary algorithms techniques, we designed primer sets specific to variants B.1.1.7, B.1.351, P.1 and respectively. Starting from sequences openly available in the GISAID repository, our pipeline was able to deliver the primer sets for each variant. In-silico tests show that the sequences in the primer sets present high accuracy and are based on 2 mutations or more. In addition, we present an analysis of key mutations for SARS-CoV-2 variants. Finally, we tested the designed primers for B.1.1.7 using RT-PCR. The presented methodology can be exploited to swiftly obtain primer sets for each new variant, that can later be a part of a multiplexed approach for the initial diagnosis of COVID-19 patients.

## Introduction

SARS-CoV-2 mutates with an average evolutionary rate of 10^−4^ nucleotide substitutions per site each year^1^. As the pandemic of SARS-CoV-2 continues to affects the globe, researchers and public health officials constantly monitor the virus for variants of concern (VOC) with acquired mutations that may pose a treat to global health such as: a higher rate of transmissibility, change in epidemiology, virulence, clinical presentation, mortality, vaccine/therapeutics resistance, or decrease in effectiveness of public health measures^2^.

On December 14th, 2020, Public Health authorities in England reported a new SARS-CoV-2 variant, ^3–5^, which belongs to the B.1.1.7 (which include all Q. lineages used for fine geographical localization of the variant) Pango lineage^6,7^, GRY clade from GISAID (Global Initiative on Sharing All Influenza Data)^3,8,9^, Nextstrain clade 20I(V1)^10^. This was the first VOC raised by the World Health Organization (WHO), and was recently termed as the Alpha variant^2^. This variant presents 14 non-synonymous mutations, 6 synonymous mutations, and 3 deletions. The multiple mutations present in the viral RNA encoding for the spike protein (S) are of most concern, such as the deletion Δ69-70 (Δ21765-21770), deletion Δ144 (Δ21991-21993), N501Y (A23063T), A570D (C23271A), P681H (C23604A), D614G (A23403G), T716I (C23709T), S982A (T24506G), D1118H (G24914C)^4,9^. Additional amino acid changes in the Spike domain, found only in a subset of the population, which are being monitored by the WHO, include those in the 484K and 452R positions^2^. The SARS-CoV-2 S protein mutation N501Y alters the protein interactions involved in receptor binding domain. The N501Y mutation has been shown to enhance affinity with the host cells ACE2 receptor^9,11^ and to be more infectious in mice^12^. The mutation P681H is adjacent to the furin cleave site of the S protein and, although there is evidence that it might play a role in SARS-CoV-2 transmission and infection ^13–15^, the effect of such mutations is still under debate^16,17^. The D614G mutation has been found to enhances viral replication, viron density and infectivity^18,19^. The alpha variant has 43 to 90% higher reproduction number than pre-existing variants increasing its transmissibility ^3,20,21^. The jury is still out on its effect on clinical severity and outcome, although there is evidence of increased clinical severity^22^ and increased mortality risk when compared with previous variants^23^. The Alpha variant remains susceptible to most monoclonal antibody therapy treatment targeting Spike protein for Covid-19 like: Bamlanivimab-etesevimab, Casirivimab-imdevimab, and others ^24–28^. Plus it has minimal impact on neutralization by convalescent and post-vaccination sera^29–32^. The presence of the Alpha variant rapidly increasing in the UK earlier this year, spreading across the globe becoming the first major global circulating VOC ^8,9,21,33,34^, before the arrival of the most recent VOC, the Delta variant which is now displacing other variants^8,35–37^.

In parallel, on December 18th, 2020, the WHO declared a new VOC, the Beta variant, first identified by the South African authorities that was rapidly spreading across 3 of their provinces and displacing other variants^5,38^. This variant is also knows know as B.1.351 Pango lineage, GH/501Y.V2 GISAID clade, and 20H (V2) Nextrain clade. The B.1.351 variant harbours 19 mutations, with 9 of them situated in the Spike protein: mutations N501Y (A23063T), E484K (G23012A), and K417N (G22813T) are at key residues in the receptor-binding domain (RBD) domain of S protein, L18F (C21614CT), D80A (A21801C), and D215G (A22206AG) are in the N-terminal domain, A701V (C23664T) in loop 2, and D614G (A23403G)^38^. Additional amino acid changes being monitored by the WHO are those on L18F position^2^. Although the B.1.351 variant also has the N501Y mutation in the Spike protein, similarly to the B.1.1.7 variant in the UK, the B.1.351 variant arose independently in a different SARS-CoV-2 lineage, which forms part of the 20H/501Y.V2 phylogenetic clade in Nextstrain^10^. Although uncommon in SARS-CoV-2 variant strains, the E484K mutation has been shown to moderately enhance binding affinity of the ACE2 receptor^11^. Mutation K417N is located in a key contact between the S protein and the human ACE2 receptor: a previous mutation in this residue has been shown to contribute to enhanced affinity of SARS-CoV-2 to the ACE2 receptor when compared to SARS-CoV^39,40^. The E484K and K417N mutations in combination with the N501Y have been shown to affect neutralization by monoclonal antibodies and convalescent sera^38,41–46^. As such the Beta variant has reduced susceptibility to monoclonal antibody therapy like bamlanivimab and etesevimab^25,28,47,48^ and it also shows reduced neutralization to convalescent and post-vaccination sera^27,31,32,49^, however it remains susceptible to casirivimab and imdevimab monoclonal antibody therapies^25,28,50^. The Beta variant also has a 50% increase in transmission rate^51^, and is associated with an increase in excess deaths per week^38^. The Beta variant spread across Africa becoming the main VOC in this region earlier this year^6,8,34^, before being displaced by the Delta variant.

On early January, Japan reported a new SARS-CoV-2 variant from 4 travellers from Brazil^52^. This variant was later reported to be widely circulating in Brazil^33,53^. On the 11th of January the WHO assigned it as VOC, the Gamma variant^2^, it is also know as P.1 Pango lineage, GR/501Y.V3 GISAID clade or 20J (V3) Nexstrain clade. The P.1 variant harbours 17 unique amino acid changes, three deletions, four synonymous mutations, and one 4-nucleotide insertion. The P.1 Variant harbours similar mutations in the Spike protein to those find in the B.1.1.7 and B.1.351 variant like the N501Y, E484K, K417T and D614G. Additional amino acid changes being monitored by the WHO are those on 681H position ^2^. Although the rate of infection and mortality of the Gamma variant are still being investigated, studies have shown a change in pattern of infection, as well as a significant change in case fatality rates associated with this variant suggesting a change in pathogenicity and virulence profile of the Gamma variant^54,55^. The Gamma variant significantly reduces the susceptibility to bamlanivimab and etesevimab monoclonal antibodies^28,47,48^, but remains susceptible to casirivimab and imdevimab monoclonal antibody therapies^28,50^. The Gamma variant is more resistant to neutralization by convalescent and post-vaccination sera^56^, plus an increase in reinfection cases has been associated to this variant^57–59^, raising concern about the need to test monoclonal antibodies in clinical against new VOC, and plausible updates to mRNA vaccines to avoid a potential loss of clinical efficacy^60^. The P.1 variant is effects still requires further study, it has been identified in the Amazonia in Brazil, as in other countries^10,53,57^. The Gamma variant spread across South America becoming the main VOC in this region until this day^8^.

On May 11^*th*^ 2021, the WHO announced a VOC, the Delta variant, which was first identified in India and linked to the rapid and deathly second wave of the country^33,61^. The Delta variant is also knows as B.1.617.2 Pango lineage (which include all AY. lineages used for fine geographical localization of the variant and do not imply any functional biological differences)^37^, G/478K.V1 GISAID clade or 21A Nexstrain clade. The B.1.617.2 harbours the next mutations in the Spike protein T19R, *l::*. 156-157, L452R, T478K, D614G, P681R, ΔR158G and D950N^33,61–64^. Additional amino acid changes being monitored by the WHO are those on 417N position ^2^. The L452R mutation has been shown to increase SARS-CoV-2 viral infectiousness and replication^65^. Interestingly. L452R and E484Q have been found to disrupt the interfacial interactions of the Spike RBD with neutralizing antibodies^66,67^. Furthermore, the L452R has been found to be a positive adaptive mutation driving the spread of similar variants with these mutations such like the ones in California^68^. The T478K is located at the interface of the Spike/ACE2 interaction domain^15,69^, and enhance stabilization of the Spike RBD with the ACE2 complex^70^. The Delta variant has been show to lead to a 64% higher household transmission rate^71^, higher viral loads^72,73^ and increase in hospital and ICU admissions when compared with the Alpha variant^74,75^. It has been associated to an increase in infections and excess deaths across the globe^64,72,76,77^. The Delta variant has been shown to be resistant to monoclonal antibody therapy treatment with bamlanivimab^25,28,47,78^, but remains susceptible to casirivimab, imdevimab and etesevimab^25,28,47,50^. Furthermore, the Delta variant has reduce neutralization by convalescent^25^ and modestly reduced sensitivity to Comirnaty/BNT162b2 mRNA vaccine^75,78–80^ and ChadOx-1 vaccine sera^36,75,80^ requiring two-doses for full protection. Although several cases of reinfection from the Delta variant have been documented in partially and fully vaccinated individuals and previously Covid-19 infected patients^81^, the severity of infection and risk of hospitalization remains lower in vaccinated individuals^74,82^. Therefore, the main concern remains in the greater burden seeing so far on health-care services due to outbreaks on unvaccinated population^74^. The Delta variant has spread like wildfire across the globe rapidly since its explosive rise in India on May, it has now become the major VOC globally rapidly displacing other variants^8,35,71,73^. Given its high prevalence and clinical implications many countries have started implemented stronger vaccine policies^83^, considering the need of additional doses for vulnerable population^84^ and the need for vaccine updates to combat waning immunity^85^.

Given the rapid spread of these VOC, and their impact on global health, it has become of utmost importance for countries to be able to rapidly identify and detect them. Several diagnostic kits have been proposed and developed to diagnose SARS-CoV-2 infections. Most kits rely on the amplification of one or several genes of SARS-CoV-2 by real-time reverse transcriptase-polymerase chain reaction (RT-PCR)^86,87^. Recently, Public Health England was able to identify the Alpha variant through their national surveillance system which allowed them to notice the rise in Covid-19 positive cases in South Easter England. The Alpha variant was detected through the increase in S-gene target failure (negative results) from the otherwise positive target genes (N, ORF1ab) in the three target gene assay in their RT-PCR diagnostic tests, and random whole genome sequencing of some of this positive samples^3,67^. In the case of the Beta^38,88,89^ and Gamma^53,90^ variants were identify through the increase in positive Covid-19 cases and deaths and by random whole genome sequencing of positive Covid-19 samples, through their national and international diagnosis and surveillance system efforts. In the case of the Delta variant, most countries rely on the clinical differences between the variants and if the RT-PCR has a S fall out (indicating Alpha variant) or not indicating a plausible Delta variant^75^. Although several PCR methods and kits for the identification of this variants have been proposed^91,92^, they usually rely on the well know Spike protein mutations, which often are shared with other VOIs and variants of SARS-CoV-2, and rapidly get obsolete due to the virus evolution^93^. Making sequencing the only reliable way to diagnose it, however this is cost prohibiting and many times not available in lower income countries.

In a previous work^94^, we developed a methodology based on deep learning, able to generate a primer set specific to SARS-CoV-2 in an almost fully automated way. Then, we reduced the necessary time by half using evolutionary algorithms^95^. When compared to other primers sets suggested by GISAID, our approach proved to deliver competitive accuracy and specificity. Our results for SARS-CoV-2 detection, both *in-silico* and with patients, yielded 100% specificity, and sensitivity similar to widely-used diagnostic qPCR methods. One of the main advantages of the proposed methodology was its ease of adaptation to different viruses or mutations, given a sufficiently large number of complete viral RNA sequences. In this work, we improved the existing semi-automated methodology, making the pipeline completely automated, and created primer sets specific for the SARS-CoV-2 variants B.1.1.7, B.1.351, P.1, B.1.6173.*, and B.1.1.519 in under 10 hours for each case study. The developed primer sets, tested *in-silico*, proved not only to be specific, but also to be able to distinguish between the different variants. In addition, we have validated our B.1.1.7 primers by RT-PCR, finding them to be specific to B.1.1.7 and sensitive enough for detection by this method. With this new result, we believe that our method represents a rapid and effective diagnostic tool, able to support medical experts both during the current pandemic, as new variants of SARS-CoV-2 may emerge, and possibly during future ones, as still unknown virus strains might surface.

## Results

### Variant B.1.1.7

As explained in the Methods section, the first step is to run a Convolution Neural Network (CNN) classifier on the data. This yields a classification accuracy of 99.66%. Secondly, from an analysis of the features constructed by the CNN, we extract 7,127 features, corresponding to 21-bps sequences. Next, we run a state-of-the-art stochastic feature selection algorithm 10 times, to uncover the most meaningful ‘21-bps’ features for the identification of variant B.1.1.7. While the best result corresponds to a set of 16 ‘21-bps’ features, using only one is enough to obtain over 99% accuracy.

These features represent good candidates for forward primers. From the 10 runs, we get 10 different ‘21-bps’ features: 5 out of the 10 point to mutation Q27stop (C27972T), 3 point to mutation I2230T (T6954C), and 2 to a synonymous mutation (T16176C). Using Primer3Plus we compute a primer set for each of the 10 features, using sequence EPI_ISL_601443 as the reference sequence. Only the two ‘21-bps’ features that include mutation T16176C are suitable for a forward primer. The two features are **ACCTCAAGGTATTGGGAACCT** and **CACCTCAAGGTATTGGGAACC**: it is easy to notice that the two features are actually part of the same sequence, just displaced by a bps, and therefore generate the same reverse primer **CATCACAACCTGGAGCATTG**.

For further analysis, we check the presence of the signature mutations of B.1.1.7; T1001I, A1708D, I2230T, SGF 3675-3677 deletion, HV 69-70 deletion, Y144 deletion, N501Y, A570D, P681H, T716I, S982A, D1118H, Q27stop, R52I, Y73C, D3L and S235F of variant B.1.1.7^96^. To verify the presence of mutations, we generate 21-bps sequences, with 10 bps before and after the mutation, and search for their presence: e.g., mutation N501Y (A23063T) corresponds to sequence **CCAACCCACT T ATGGTGTTGG**.

Using the generated ‘21-bps’ sequences for the mutations, we can also test them as forward primers using Primer3Plus, which yields **TGATATCCTTGCACGTCTTGA** in spike gene S982A as the only possible forward primer candidate, this sequence can be used for multiplex testing (Fig. 1).

**Figure 1.**
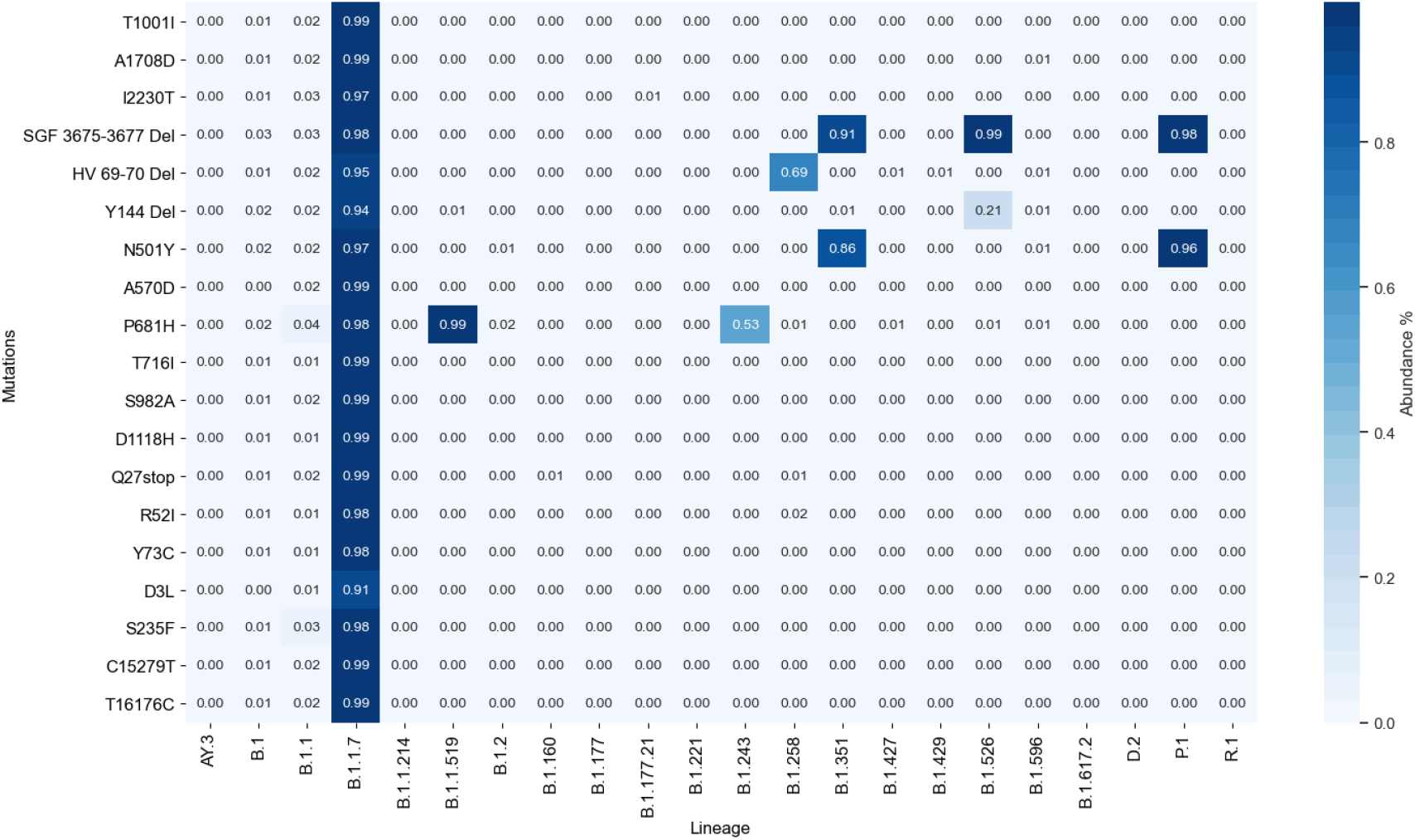
Frequency of appearance of characteristic mutations of B.1.1.7 Variant in 2,096,390 SARS-CoV-2 sequences downloaded from GISAID in August 11^*th*^, 2021.

### Variant B.1.351

Again, the first step is to run a CNN classifier on the data. This yields a classification accuracy of 99.88%. Next, we translate the CNN weights into 6,069 features, each representing a 21-bps sequence. we run the feature selection algorithm 10 times, which results in only one meaningful feature achieving over 99% accuracy. The 10 runs point to the same mutation, K417N, with 6 different sequences but only one acceptable as a primer candidate. The resulting sequence is **CTCCAGGGCAAACTGGAAATA**, with a reverse primer **TGCTACCGGCCTGATAGATT**.

We check whether 21-bps sequences containing the signature mutations T265I, K1655N, K3353R, SGF 3675-3677 deletion, L18F, D80A, D215G, R246I, K417N, E484K, N501Y, A701V, 242-244 del, Q57H, S171L, P71L, and T205I of variant B.1.351^96^ could function as a forward primer. From the Primer3Plus results using EPI_ISL_678597, mutations D215G (**TTAGTGCGTGGTCTCCCTCAG**) and Q57H (**CTGTTTTTCATAGCGCTTCCA**) can be used as forwards primers (Fig. 2).

**Figure 2.**
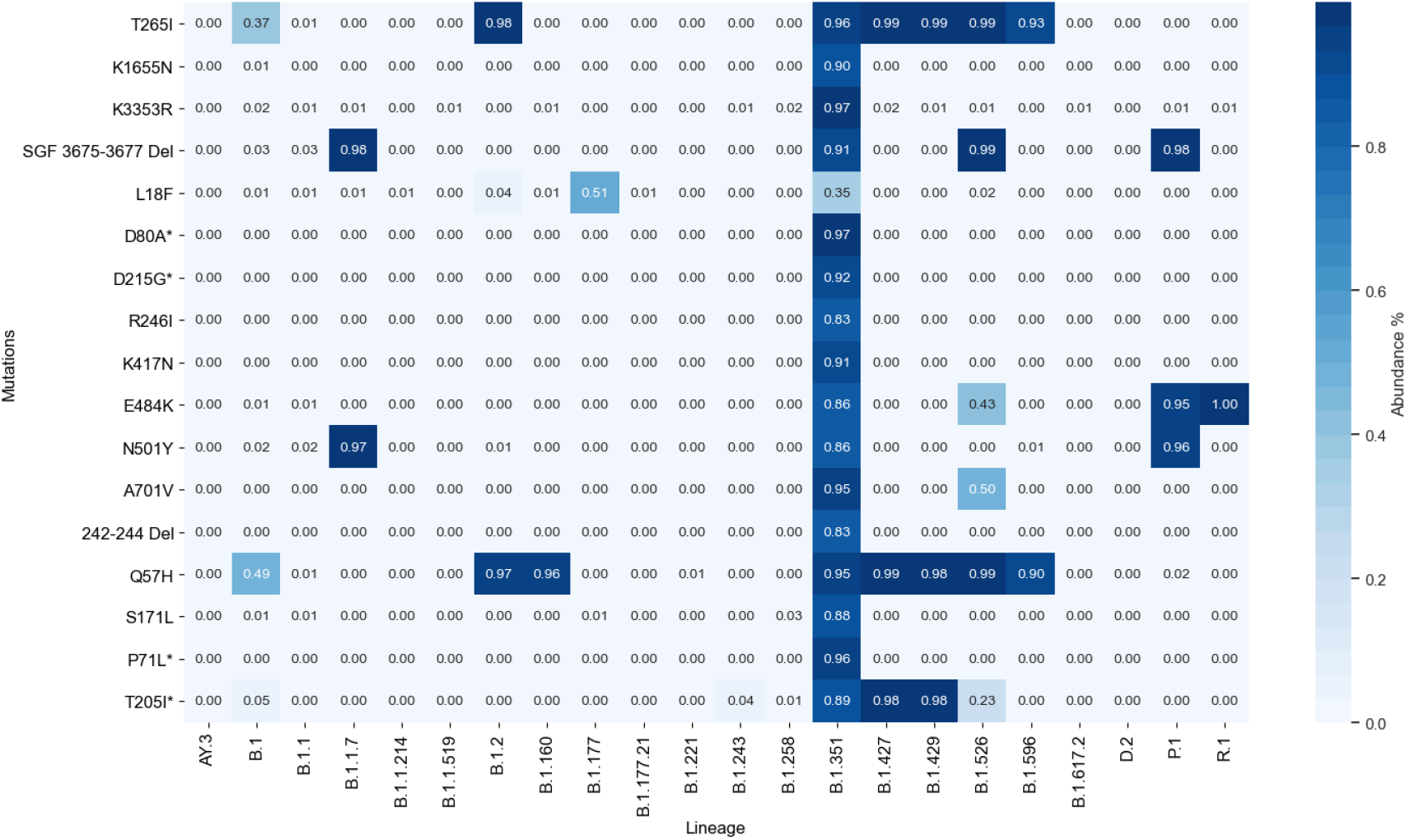
Frequency of appearance of characteristic mutations of B.1.351 Variant in 2,096,390 SARS-CoV-2 sequences downloaded from GISAID in August 11^*th*^, 2021.

### Variant P.1

The CNN classifier on the data yields a classification accuracy of 100%, in the 28 samples. Next, we translate the CNN weights into 727 features, each representing a 21-bps sequence. We run the feature selection algorithm 10 times, which results in only one meaningful feature achieving 100% accuracy. The 10 runs point to the same sequence, around the synonymous mutation T733C **ACTGATCCTTATGAAGACTTT**. The sequence cannot be used as a primer, as the Tm is too low. To compensate, we displaced the sequence two bps to the left and added a bps to rise the Tm. This procedure gave us sequence **GGCACTGATCCTTATGAAGACT**, of size 22 bps, with reverse primer **TTCGGACAAAGTGCATGAAG**.

We check whether the sequences containing signature mutations S1188L, K1795Q, E5665D, SGF 3675-3677 deletion, T20N/L18F, P26S, D138Y, R190S, K417T, EA84K, N501Y, H655Y, T1027I, E92K, ins28269-28273 AACA, and P80R of variant P.1^53^ could function as a forward primers. We put mutation T20N and L18F into the same ‘21-bps’ sequence, given their proximity. From the Primer3Plus results using EPI_ISL_792683, 3 of the generated sequences using mutations can be forward primers candidates; K417T (**CAAACTGGAACGATTGCTGAT**), H665Y (**AGGGGCTGAATATGTCAACAA**) and P80R (**AATAGCAGTCGAGATGACCAA**) (Fig. 3).

**Figure 3.**
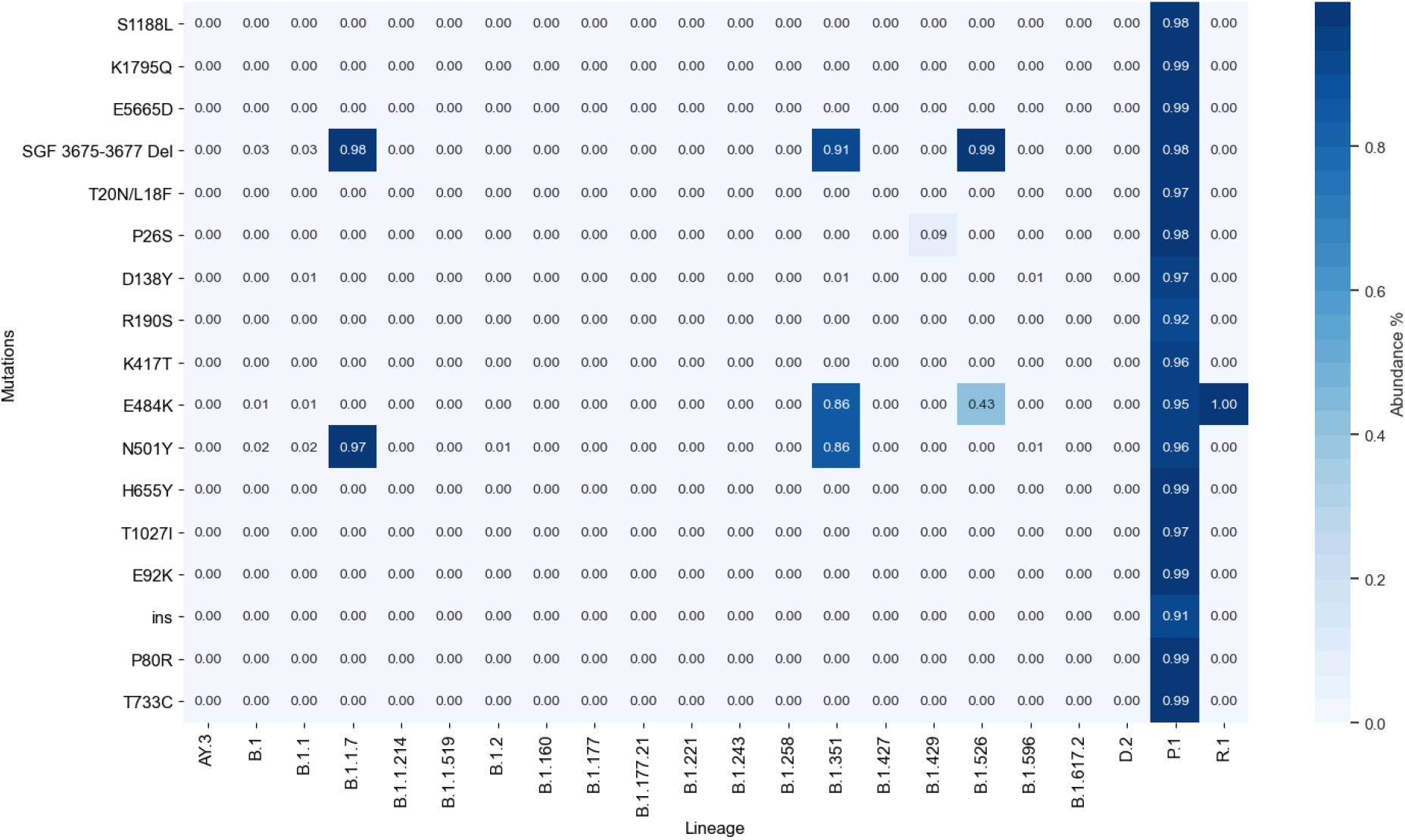
Frequency of appearance of characteristic mutations of P.1 Variant in 2,096,390 SARS-CoV-2 sequences downloaded from GISAID in August 11^*th*^, 2021.

### Variant B.1.617.2

Using the sequences of variant B.1.617.2 downloaded from the GISAID repository we ran an evolutionary algorithm 10 times to find the most important sequences to separate the B.1.617.2 from the rest of the sequences. From the results 4 out of 10 point to mutation D63G, 4 to mutation I82T and 2 to mutation T120I. The forward primer for mutation I82T is (**CTACCGCAATGGCTTGTCTT**) and for mutation D63G is (**ATGGCAAGGAAGGCCTTAAA**) using sequence EPI_ISL_1337507. Then, we make an analysis of the characteristic mutations of lineage B.1.617.2; T19R, L452R, T478K, P681R, D950N, S26L, I82T, V82A T120I, D63G, R203M, D377Y and Del 156-157^97^. Where sequences of mutations T478K (**GCCGGTAGCAAACCTTGTAAT**), V82A (**GCCAGATCAGCTTCACCTAAA**) and R203M (**GGCAGCAGTATGGGAACTTCT**) can be used as forward primers in multiplex approach (Fig. 4). It is important to consider that lineage AY.3 with B.1.617.2 are part of the *Delta* variant.

**Figure 4.**
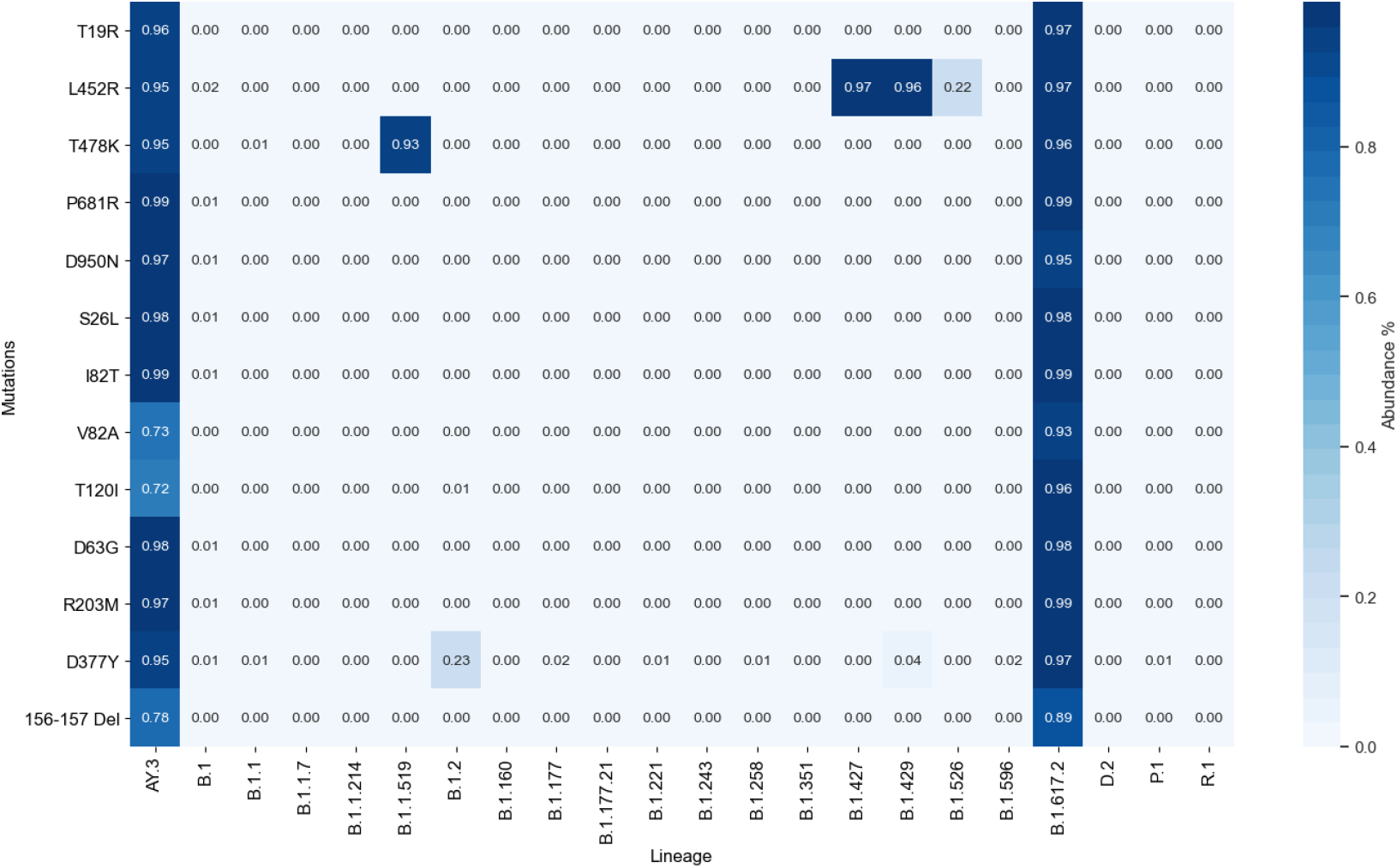
Frequency of appearance of characteristic mutations of B.1.617.2 Variant in 2,096,390 SARS-CoV-2 sequences downloaded from GISAID in August 11^*th*^, 2021.

### Experimental evaluation of B.1.1.7 Primer specificity

The RT-PCR amplification curves obtained with B.1.1.7 specific primer set B.1.1.7-1 and SARS-CoV-2 generic primers IP2 and IP4 (following the Pasteur Institute Protocol)^98^ on two SARS-CoV-2 strains are shown result in Fig. 5. As expected, only the B.1.1.7 strain is amplified the B.1.1.7-1 primers, while both the Wuhan reference strain and the B.1.1.7 strain are detected by the generic primers. Fig. 5. Comparison between non-specific primer set IP2 and IP4 and our designed primer set B.1.1.7-1 for B.1.1.7 variant and the original Wuhan SARS-CoV-2 strain.

**Figure 5.**
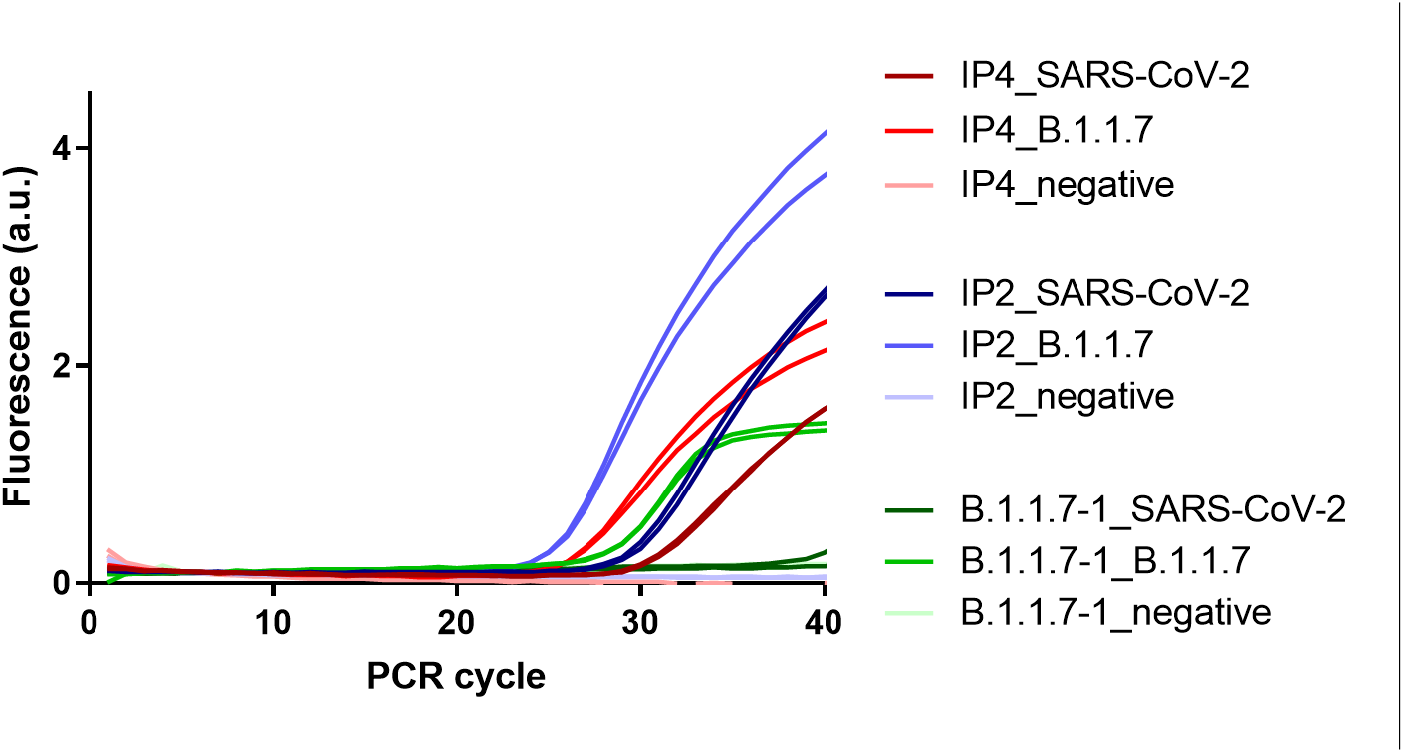
Comparison between non-specific primer set IP2 and IP4 and our designed primer set B.1.1.7-1 for B.1.1.7 variant and others.

## Discussion

A single-nucleotide mutation may not be enough to work as a specific primer for detecting SARS-CoV-2 variants. Thus, from the analysis of the characteristics mutations for each variant and the results of our pipeline, we created a list of primer sets based upon 2 or more mutation for each variant. For variant B.1.1.7 we created 3 primer sets, for variants B.1.351, and P.1 we created 2 options for each one, and for B.1.617.2 we created 3 different combinations (Table 1). Using Primer3Plus for in-silico simulations, we tried to maintain acceptable levels of temperature and an 18-22 bps size. Nevertheless, it was not always possible e.g. for P.1 variant the forward primers contain sequences size 25 bps. Another thing to consider is the product size, depending of which primer is going to be used the size will vary, as each primer falls into at least one mutation. The frequency of appearance for the different sequences is in Fig. 6.

**Table 1.**
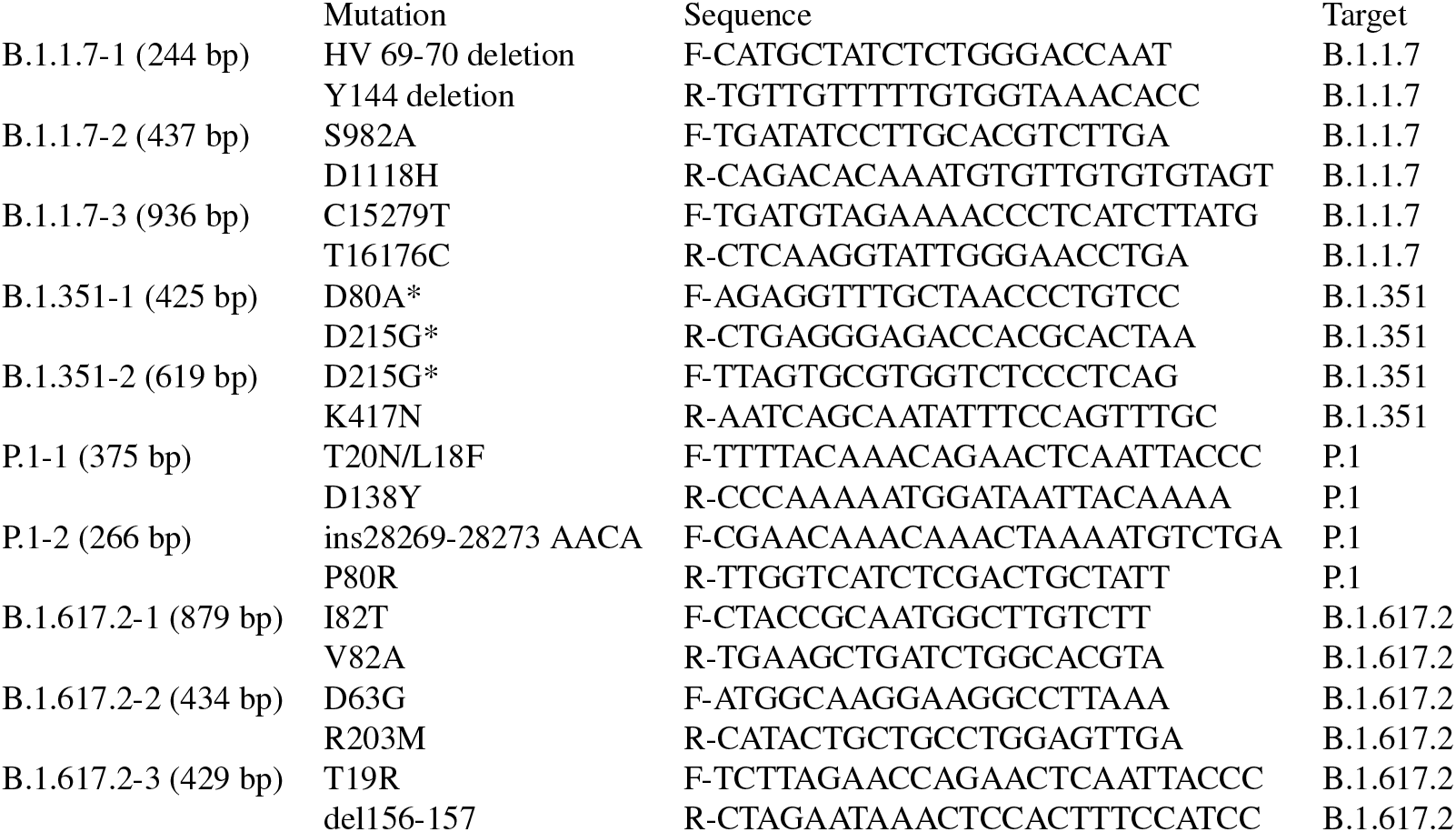
Sequences of the primer sets for B.1.1.7, B.1.351, P.1 and B.1.617.2 lineages.

**Figure 6.**
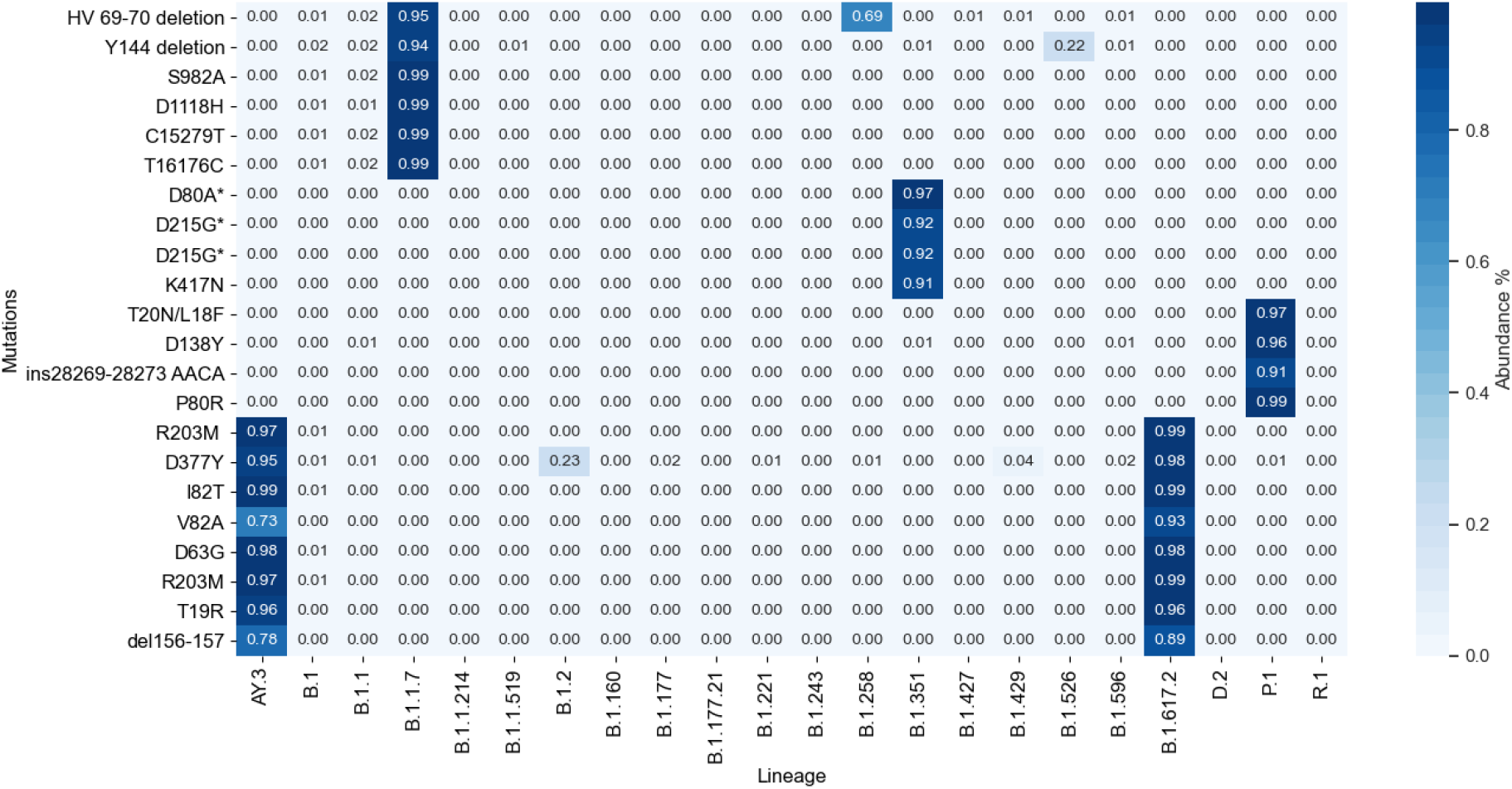
Frequency of appearance of the different primer sets in 2,096,390 SARS-CoV-2 sequences downloaded from GISAID in August 11^*th*^, 2021.

A wide variety of diagnostic tests have been used by high-throughput national testing systems around the world, to monitor the SARS-CoV-2 infection^86^. The arising prevalence of new SARS-CoV-2 variants such as B.1.1.7, B.1.351, P.1 and B.1.617.2 have become of great concern, as most RT-PCR tests to date are not be able to distinguish these new variants, not being designed for such a purpose. Therefore, public health officials most rely on their current testing systems and whole viral RNA sequencing results to draw conclusions on the prevalence of new variants in their territories. An example of such case has been seen in the UK, where the increase of the B.1.1.7 SARS-CoV-2 variant infection in their population was identified only through an increase in the S-gene target failure in their three target gene assay (N+, ORF1ab+, S-), coupled with sequencing of the virus and RT-PCR amplicons products^3^. Researchers believe that the S-gene target failure occurs due to the failure of one of the RT-PCR probes to bind, as a result of the 69-70 deletion in the SARS-CoV-2 spike protein, present in B.1.1.7^3^. This 69-70 deletion, which affects its N-terminal domain, has been occurring in several different SARS-CoV-2 variants around the world^10,99^ and has been associated with other spike protein receptor binding domain changes^9^. Due to the likeliness of mutations in the S-gene, assays relying solely on its detection are not recommended, and a multiplex approach is required^86,87,100^. This is consistent with other existing primer designs like CoV2R-3 in the S-gene^101^, that will also yield negative results for the B.1.1.7 variant, as the reverse primer sequence is in the region of mutation P681H. A more in-depth analysis of S-dropout positive results can be found in Kidd et al.^102^.

Given the concern for the increase in prevalence of the new variants SARS-CoV2 B.1.1.7, B.1.351, P.1 and B.1.617.2 and their possible clinical implication in the ongoing pandemic, diagnosing and monitoring the prevalence of such variants in the general population will be of critical importance to help fight the pandemic and develop new policies. In this work, we propose possible primer sets that can be used to specifically identify the B.1.1.7, B.1.351, P.1 and B.1.617.2 SARS-CoV2 variants. In addition, we tested the detection and specificity for the variant B.1.1.7. We believe that all the proposed primer sets can be employed in a multiplexed approach in the initial diagnosis of Covid-19 patients, or used as a second step of diagnosis in cases already verified positive to SARS-CoV-2, to identify individuals carrying the B.1.1.7, B.1.351, P.1 or B.1.617.2 variant. In this way, health authorities could better evaluate the medical outcomes of this patients, and adapt or inform new policies that could help curve the rise of variants of interest. Although the rest of the proposed primer sets delivered by our automated methodology will still require laboratory testing to be validated, our methodology can enable the timely, rapid, and low-cost operations needed for the design of new primer sets to accurately diagnose new emerging SARS-CoV-2 variants and other infectious diseases.

## Data and Methods

### Methods

For the discovery of the regions of interest in each variant we used two methodologies, that yield similar results. For the variants B.1.1.7, B.1.351 and P.1 we used a methodology based in Convolution Neural Networks (CNN) as explained in^94^. For the variants B.1.617.2 we used a methodology based in Evolutionary Algorithms (EAs)^95^.

#### CNN

Following the procedure described in Lopez et al.^94^, there are 4 steps for the automated design of a specific primer for a virus: (i) run a CNN for the classification of the target virus against other strains, (ii) translate the CNN weights into 21-bps features, (iii) perform feature selection to identify the most promising features, and (iv) carry out a primer simulation with Primer3Plus^103^ for the features uncovered in the previous step. While in^94^ the proposed pipeline was only partially automatic, and still required human interventions between steps, in this work all steps have been automatized, and the whole pipeline has been run with no human interaction. The experiments, from downloading the sequences to the final *in-silico* testing of the primers, take around 16 hours of computational time on a standard end-user laptop, for each variant considered.

In a first step, we train a convolution neural network (CNN) using the training and testing sequences obtained from GISAID. The architecture of the network is shown in Fig 7, and is the same as the one previously reported in^94^. Next, if the classification accuracy of the CNN is satisfying (*>* 99%), using a subset of the training sequences we translate the CNN weights into ‘21-bps’ features, necessary to differentiate between the targeted variant samples and all the others. The length of the features is set as 21 bps, as a normal length for primers to be used in PCR tests is usually 18-22 bps. Then, we apply recursive ensemble feature selection (REFS)^104,105^ to obtain a reduced set of the most meaningful features that separate the two classes. Finally, we simulate the outcome of treating the most promising features obtained in the previous step as primers, using Primer3Plus^103^ in the canonical sequences EPI_ISL_601443 for B.1.1.7^3^, EPI_ISL_678597 for B.1.351 and EPI_ISL_804814 for P.1^53^.

**Figure 7.**
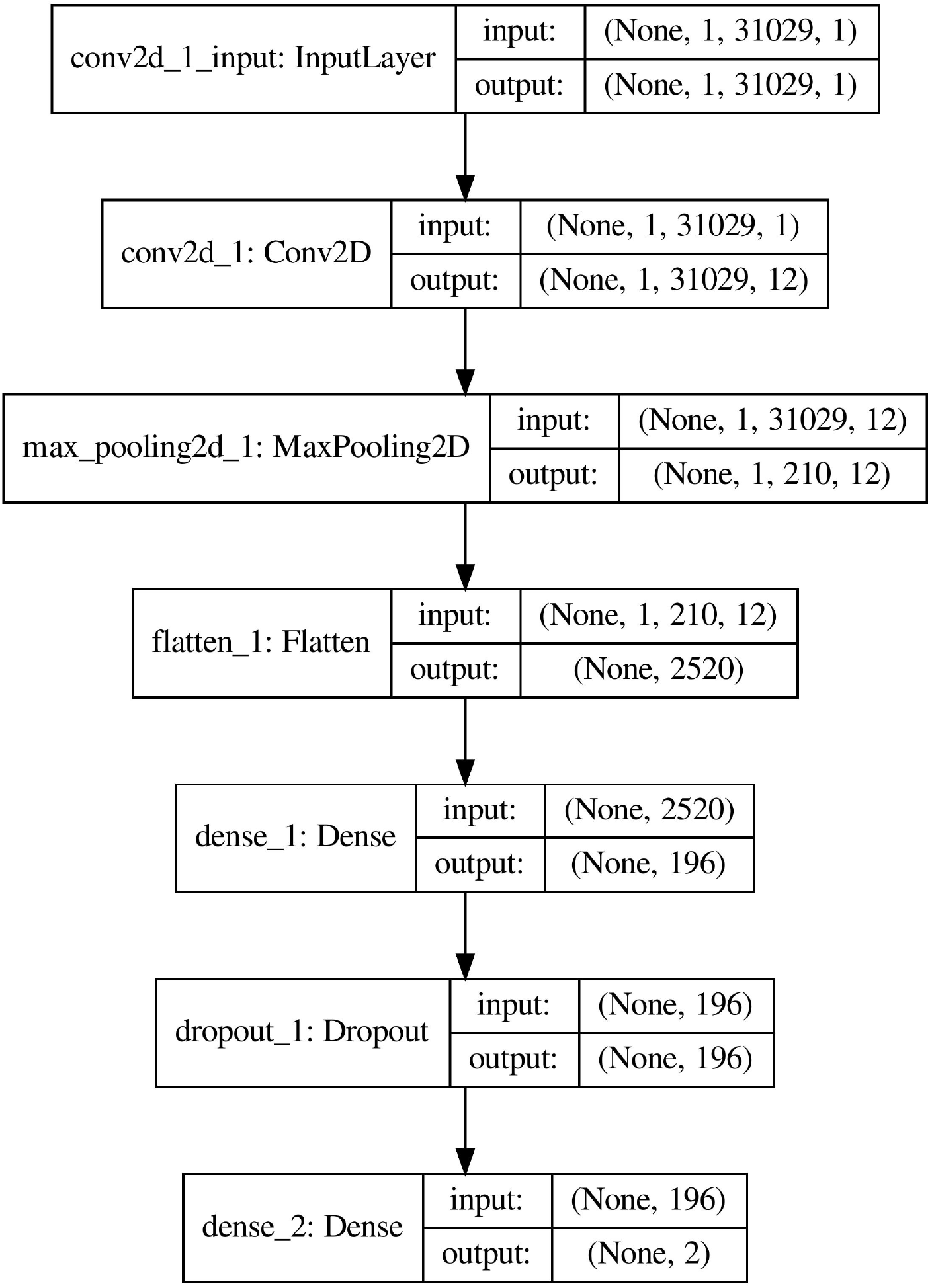
CNN architecture used to classify variants B.1.1.7, B.1.351, and P.1.

#### Evolutionary Algorithms

Another approach to generating primers is to use EAs. This approach has the advantage that we can parallelize the procedure and reduce the time required to get the regions of interest in comparison to the CNN-based method. In comparison to the 16 hours required to the CNN approach, each single run lasts around 62 minutes with 5 threads on a 64-bit Windows 10 laptop with Intel Xeon E-2186M. For the EA, We create a set of individuals of size 21-bps randomly considering the available samples of the variant of interest (e.g. B.1.1.519) and other variants. Next, we calculate what is known as cost function, which is given by the following:

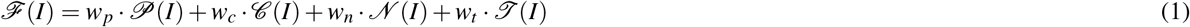

with *w*_*p*_, *w*_*c*_, *w*_*n*_, *w*_*t*_ representing the weights associated to each term.

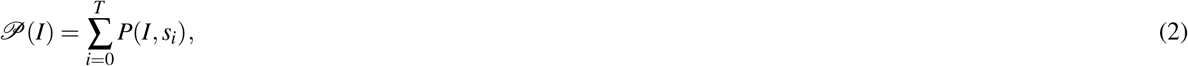

𝒫 (*I*) is evaluating the presence of the sequence selected as candidate primer *I* inside training samples labeled with the variant of interest, and its absence from samples of other variants, *T* is the number of samples in the training set, *s*_*i*_ is the *i*-th sample in the training set. Function *P* is defined as:

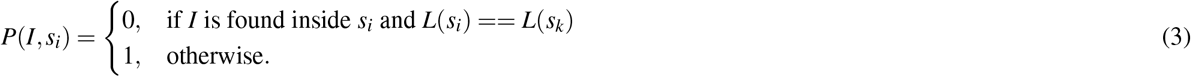

where *L*(*s*) returns the class label of sample *s*. In other words, *P*(*I, s*_*i*_) equals 1 if sequence *I* is found inside a sample with the same class label as sample *s*_*k*_, the origin of sequence *I*. So, if the 21-bps sequence *I* is found inside a sample that does not belong to the variant of interest, or is not found in a sample that belongs to the variant of interest, the solution is penalized.

The second term of the weighted sum takes into account the GC content of the candidate primer:

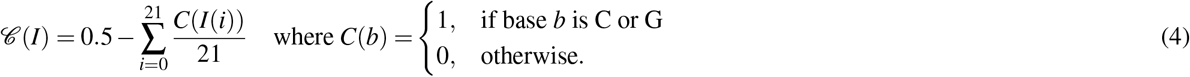

where *I*(*i*) represents the base in position *i* inside sequence *I*. The following element of the weighted sum is *𝒩*, defined as:

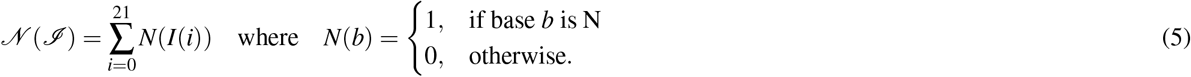

that takes into account the presence of N symbols in the sequence, indicating an error in the read. The ideal primer candidate should only contain A, C, G, or T values.

The final term tackles the requirement of having a melting temperature *T*_*m*_ centered around 60^*?*^. Specialized literature^103^ provides an equation to compute *T*_*m*_ for a sequence *I*:

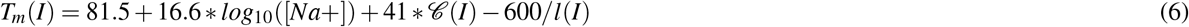

where *𝒞* (*I*) is the content of C and G bases in sequence *I*, as described in Equation 4, [*Na*+] is the molar sodium concentration, and *l*(*I*) is the length of sequence *I*, in bps. We used the value of [*Na*+] = 0.2 as described in^103^, while *l*(*I*) = 21 by design. The term taking into account *T*_*m*_ will then be:

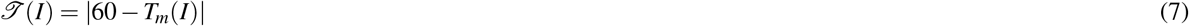

The EA is set with a population of size *µ* = 200, generating offspring of size *λ* = 200. The entire population is replaced by its offspring at each generation, using a (*λ, µ*) replacement strategy, with a tournament selection of size *τ* = 2, a mutation acting on integer values, a one-point crossover, and a stop condition set on 100 generations. For more details, please refer to^95^.

#### Experimental evaluation of B.1.1.7 Primers

Amplification efficiency of the designed primer sets were evaluated using viral RNA extracts from two sequenced SARS-CoV-2 strains: the original Wuhan strain 210207 (GISAID N° EPI_ISL_437689) and VOC B.1.1.7 strain (GISAID N° EPI_ISL_683466). Viral RNA were extracted from infected cell culture supernatants using the NucleoSpin Dx Virus kit (Macherey-Nagel), following the manufacturers’ protocol. Viral RNA extracts (5 µL) were analyzed either using the IP2/IP4 dualplex real-time reverse-transcriptase (RT)–PCR assay, developed by following the Pasteur Institute and targeting conserved regions of the SARS-CoV-2 RdRP gene^98^, or primer set B.1.1.7-1 (Table 1), using the LightCycler EvoScript RNA SYBR Green I Master kit (Roche). Both RT-PCR assays were conducted on a LightCycler® 480 System (Roche), using the thermal cycling program described in the Pasteur Institute protocol^98^.

### Data

#### Variant B.1.1.7

From the GISAID repository we downloaded 10,712 SARS-CoV-2 sequences on December 23, 2020. After removing repeated sequences, we obtained a total of 2,104 sequences labeled as B.1.1.7, and 6,819 sequences from other variants, for a total of 8,923 samples. B.1.1.7 variant samples were assigned label 1, and the rest were assigned label 0 for the CNN discovery described in detail in^94^. Next, from the found combinations and known mutations we generated primer sets and test them in 2,096,390 SARS-CoV-2 sequences downloaded in August 11^*th*^, 2021, where 1,051,740 sequences are B.1.1.7. The total number of sequences by lineage is in the Table 2 on the supplemenatary material.

**Table 2.** Number of sequences by lineage downloaded from the GISAID repository on August 11^*t*^*h*, 2021.

| Lineage | Number of Sequences |
| --- | --- |
| AY.3 | 17504 |
| B.1 | 82953 |
| B.1.1 | 47373 |
| B.1.1.7 | 1051740 |
| B.1.1.214 | 18066 |
| B.1.1.519 | 23376 |
| B.1.2 | 101442 |
| B.1.160 | 27353 |
| B.1.177 | 74475 |
| B.1.177.21 | 13048 |
| B.1.221 | 13510 |
| B.1.243 | 12841 |
| B.1.258 | 14061 |
| B.1.351 | 30650 |
| B.1.427 | 18239 |
| B.1.429 | 39345 |
| B.1.526 | 49786 |
| B.1.596 | 10939 |
| B.1.617.2 | 366831 |
| D.2 | 12772 |
| P.1 | 59692 |
| R.1 | 10394 |
| Total | 2096390 |

#### Variant B.1.351

From the GISAID repository, we downloaded 326 sequences of the B.1.351 variant on January 7, 2021. We added the 326 sequences to the 8,923-sample dataset from the B.1.1.7 experiment, obtaining a total of 9,249 sequences, where we assigned the label 1 for sequences belonging to variant B.1.351, and 0 to the rest of the samples using CNN to find the primers. Next, we generated primer sets and test them in 2,096,390 SARS-CoV-2 sequences downloaded in August 11^*th*^, 2021 where 30,650 sequences are B.1.351.

#### Variant P.1

From the GISAID repository, we downloaded 28 non-repeated sequences of the P.1 variant on January 19, 2021. We added the 28 sequences to the 8,323 sequences of several other variants, including B.1.1.7 and B.1.351, for a total of 8,351 sequences. We assigned label 1 to sequences belonging to variant P.1, and 0 to the rest of the samples using CNN to find the primers. Next, we generated primer sets and test them in 2,096,390 SARS-CoV-2 sequences downloaded in August 11^*th*^, 2021 where 59,692 sequences are P.1.

#### Variant B.1.617.2

From the GISAID repository, we downloaded 836 sequences of the B.1.617.2 variant on May 5th, 2021. We added sequences to 6,819 sequences of other variants, and we assigned label 1 to sequences belonging to variant B.1.617.2, and 0 to the rest of the samples using EAs to find the primers. Finally, we generated primer sets and test them in 2,096,390 SARS-CoV-2 sequences downloaded in August 11^*th*^, 2021 where 366,831 sequences are B.1.617.2.

## Additional information

The authors declare no competing interests.

## Author contributions statement

CAP, LMM, made the biological analysis, and primer design. ALR and AT made the programming, data collection, and experiments in silico. EC, ADK and JG made the experiment and study design. EC and PT made the laboratory testing. CAP and ALR wrote the the article, all authors contributed to editing of the article.

## Supplementary Material

